# Bacterial community changes with cryoconite granule size and their susceptibility to exogenous nutrients on 10 glaciers in northwestern Greenland

**DOI:** 10.1101/514083

**Authors:** Jun Uetake, Naoko Nagatuska, Yukihiko Onuma, Nozomu Takeuchi, Hideaki Motoyama, Teruo Aoki

## Abstract

Cryoconite granules, which are dark-colored biological aggregates on glaciers, effectively accelerate the melting of glacier ice. Bacterial community varies with granule size, however, community change in space and their susceptibility to environmental factors has not been described yet. Therefore, we focused on bacterial community from 4 different granule sizes (30-249 μm, 250- 750 μm, 750-1599 μm, more than 1600 μm diameter) in 10 glaciers in northwestern Greenland and their susceptibility for exogenous nutrients in cryoconite hole. A filamentous cyanobacterium *Phormidesmis priestleyi*, which has been frequently reported from glaciers in Arctic was abundant (10-26%) across any size of granules on most of glaciers. Bacterial community across glaciers became similar with size increase, and whence smallest size fractions contain more unique genera in each glacier. Multivariate analysis suggests that phosphate, which is significantly higher in one glacier (Scarlet Heart Glacier), is primary associated with bacterial beta diversity. Correlation coefficients between abundance of major genera and nutrients largely changed with granule size, suggesting that nutrients susceptibility to genera changes with growth process of granule (e.g. *P. priestleyi* was affected by nitrate in early growth stage).

## Introduction

Cryoconite granules are dark-colored spherical aggregates on glacier surface, consisting of mineral particles, organic matter and microorganisms (Gerdel and Drouet 1960, Cook et al. 2015). Their granular structure is formed by filamentous cyanobacteria and further cemented by extracellular polymeric substances produced by microorganisms (Takeuchi et al. 2001, Hodson et al. 2010, Langford et al. 2010,). Because of dark coloration of the cryoconite granules, they effectively reduce the glacier surface albedo and accelerate the melting of glacier ice (Anesio and Laybourn-Parry 2011, Takeuchi et al. 2014, Cook et al. 2015, Musilova et al. 2016). On Greenland Ice Sheet, formation of cryoconite granules has been suggested to contribute substantially to glacier mass loss (Anesio and Laybourn-Parry 2011, Takeuchi et al. 2014, Cook et al. 2015) and are estimated to become a greater impact in the future (Musilova et al. 2016).

Supraglacial microbial community is affected by various environmental factors such as climate conditions, physical features of glacier surface and fluxes of carbon and nutrients (Stibal et al. 2012, Cook et al. 2015, Anesio et al. 2017). Previous studies suggested the hydrological factors such as ice melting rate and meltwater flow are most influential to microbial community (thermal regime affecting supraglacial melt flow: Edwards et al. 2011, slope angle as index for melt flow: Stibal et al. 2012, positive degree day as index for duration of growth: Gokul et al. 2016). Furthermore, nutrients including nitrogen and phosphorus, which are usually originated from debris and/or aerosols deposited on glaciers, are also limiting factors for the microbial community on glaciers (Stibal et al. 2007). A positive relationship between nutrients and glacier microbial community was found in long-term (1-3 years) laboratory incubation experiment of cryoconite granule under illuminate condition (Musilova et al. 2016). In contrast, a short term (2 months) nitrate addition into in-situ cryoconite holes showed minimal impacts to supraglacial bacterial community structure (Cameron et al. 2017). Furthermore, another short term (1 week) laboratory experiment of phosphorus addition together with and without glucose as carbon source showed that they do not affect the bacterial cell concentration at 0.1°C (Mindl et al. 2008). These previous studies indicated that in situ nutrients would and wouldn’t affect microbial communities and this topic is still open question. Meanwhile, endogenous biogeochemical nutrient cycles such as nitrogen cycles (Telling et al. 2012, Segawa et al. 2014), phosphor utilization by phosphatase (Stibal et al. 2009) and direct substance transfer between microorganisms (Smith et al. 2016) has been found to occur within cryoconite granules, thus, microorganisms in mature granule with high density cells and substances are possibly affected by nutrients not only from exogenous (outside of granules) but also from endogenous (inside of granules) sources.

On the glacier surface, the size of cryoconite granules usually varied from μm to mm in diameter, because filamentous cyanobacteria interweave their cells to make a granule and granule size is changed by degree of microbial growth. 16S rRNA gene amplicon sequencing of bacterial community in different granule size on a Greenland glacier revealed that the communities were distinctive among the size (diameter) of the granules and their diversity largely differed between granules of larger and smaller than 250 μm in diameter (Uetake et al. 2016). However, this study was done on a single glacier and it is difficult to presume spatial variation of microbial communities with granule size changes, because microbial communities are quite different even in adjacent glaciers (Edward et al. 2011, Lutz et al. 2016).

Therefore, at first, in order to know spatial difference of bacterial changes with granule growth across glaciers, we focused on bacterial community based on 16S rRNA gene sequencing in 4 different sizes (30-249 μm, 250- 750 μm, 750-1599 μm, more than 1600 μm) of cryoconite granules collected from cryoconite holes on 10 different neighborhood glaciers of previous study site in Uetake et al. (2016). Secondary, differences of microbial communities with granule size would change susceptibility to environmental factors, because firm aggregated granule is more stable and nutrients are potentially supply from in situ nutrient cycle such as nitrogen fixation (Telling et al. 2012) and nitrification and denitrification processes (Segawa et al. 2014). Therefore, in order to understand susceptibility of exogenous nutrients (phosphors, nitrate, nitrite and ammonium) to bacteria and bacterial diversity in each granule fractions, simple correlation and multivariate analysis method were conducted for 10 glaciers.

## Materials and Methods

### Sampling and size sorting

We conducted field research on 10 glaciers near a village of Qaanaaq located in Northwestern Greenland in June- Aug 2013 (SI Fig.1, SI Table1). Sampling duration for each glacier and access are summarized in Aoki et al. (2014). Cryoconite granules were collected from bottom of cryoconite holes at three - five points per each site by using pre-cleaned plastic pipette handled with sterile new plastic gloves. The, cryoconite granules were transferred to RNAlater (MilliporeSigma) within one third of total volume (cryconite granules, residue water and RNAlater) and kept in portable freezer (SC-DF25, Twinbird, Niigata, Japan) at −30 degrees in field and transportation, then stored in deep freezer at −80 degrees in National Institute of Polar Research (NIPR) until further processes. Because replicated cryoconite granule samples from QQ are already merged into one sample, QQ are excluded from statistical comparison (t-test, SIMPER, ANOSIM and PERMANOVA).

Meltwater in cryoconite hole were collected directly with 50ml pre-cleaned plastic bottle, then filtrated by 0.45μm pore 13 μm ϕ syringe filter for ion chromatography (5040-28521, GL Science, Tokyo, Japan) into 5ml plastic tube and kept frozen as with DNA samples until analysis. All cryoconite granules were sieved through four different meshes (30 μm mesh: NRS-030, Nippon Rikagaku Kikai, Tokyo, 250 μm mesh: NMG66, 750 μm mesh: NMG27 and 1600 μm mesh: NMG14, SEMITEC, Osaka, Japan) by pouring ultrapure water (Milli-Q^®^ Advantage A10; Merck Millipore, MA, USA) from lab wash bottle and sorted into four different size categories (Size 30: 30-250 μm, Size 250: 250-749 μm, Size 750: 750-1600 μm, Size 1600: more than 1600 μm).

### DNA sequencing

Genomic DNA of sorted cryoconite granules were extracted using the FastDNA™ SPIN Kit for Soil DNA Extraction (MP Biomedicals, Santa Ana, CA). Partial 16S rRNA gene sequences including the V3 and V4 regions were amplified using the primers Bakt_341F (CCTACGGGNGGCWGCAG) and Bakt_805R (GACTACHVGGGTATCTAATCC) with Illumina overhang adaptor sequences attached to their 5′. Polymerase chain reaction (PCR) cycle is 30 cycles of denaturation at 98°C for 10 s, annealing at 52°C for 30 s, and elongation at 72°C for 1 min 30 s with initial denature at 98°C for 3 min and additional elongation 72°C for 3 min by GeneAmp PCR System 9700 (Applied Biosystems, CA, USA). Subsequent clean-up, index PCR were performed following Illumina methods for “16S metagenomic sequencing library preparation” (https://support.illumina.com/content/dam/illumina-support/documents/documentation/chemistry_documentation/16s/16s-metagenomic-library-prep-guide-15044223-b.pdf). All samples were pooled into a flow cell of MiSeq sequencer (Illumina, San Diego, CA) in NIPR and sequenced by Miseq V3 reagent kit. All sequence data are available from DDBJ Sequence Read Archive: https://www.ddbj.nig.ac.jp/dra/index-e.html (DRA007055).

### Amplicon sequence variant analysis

In order to understand much more detail sequences differences than conventional 97% operational taxonomic unit (OTU) method, we used newly developed DADA2 (1.4) method which incorporates error model and is able to infer with single nucleotide resolution (Callahan et al. 2016). All potential chimera sequences were removed by DADA2. Taxonomy was assigned by naive Bayesian classifier method in The Ribosomal Database Project (RDP) Classifier (Wang et al. 2007) implemented in DADA2 program against costumed Silva 128 database (Quast et al. 2013). Costumed Silva 128 database is including ordinary sequences and sequences from Uetake et al. 2016 which studied cryoconite granule 16S rDNA in same region. Alpha diversity and Weighted UniFrac distances analysis (Lozupone et al. 2006) were analyzed by QIIME (Caporaso et al. 2010). Potential function of bacterial communities related to nutrient cycling were estimated by “Tax4Fun” package in R using Silva 123 database (Aßhauer et al. 2015, R Core Team 2017).

#### Ion concentrations

Ion concentrations of NH_4_^+^, NO_3_^−^, PO_4_^3−^ and SiO_3_^2−^ were determined calorimetrically using an Autoanalyzer II (Bran+Luebbe, Norderstedt, Germany) in NIPR.

#### Data analysis

Difference of alpha diversity and nutrients across glaciers were analyzed by one way Analysis of variance (ANOVA) and Tukey’s Honest Significant Difference (Tukey HSD) test by “base” function of R”. Analysis of similarities (ANOSIM) and permutational multivariate analysis of variance (PERMANOVA) were performed using PRIMER 7 (Plymouth, UK). Samples from Indicator species of each sites were detected by IndVal functions of “labdsv” package (https://cran.r-project.org/web/packages/labdsv/labdsv.pdf) in R.

## Results

### 1: Taxonomy compositions in different size fractions across the 10 glaciers

Within total 38 sites across 10 glaciers, 30 sites with average 25445 sequences (SD: 7854) for size30, 30 sites with average 26302 sequences (SD: 6605) for size250, 31 sites with average 27650 sequences (SD: 7242) for size1600 and 26 sites with average 32011 sequences (SD: 8765) for size1600 were analyzed for downstream analysis. *Cyanobacteria* were most abundant in the granules of all size fractions and were followed by *Proteobacteria* or *Bacteroidetes* (Fig.1) except size 30 in SER, MJ and NT (SER: *Bacteroidetes* 30%, MJ and NT: *Proteobacteria* 23% and 23%) and size 1600 in SUN and BD (*Proteobacteria*, SUN: 24%, BD1: 29%).

**Figure 1:**
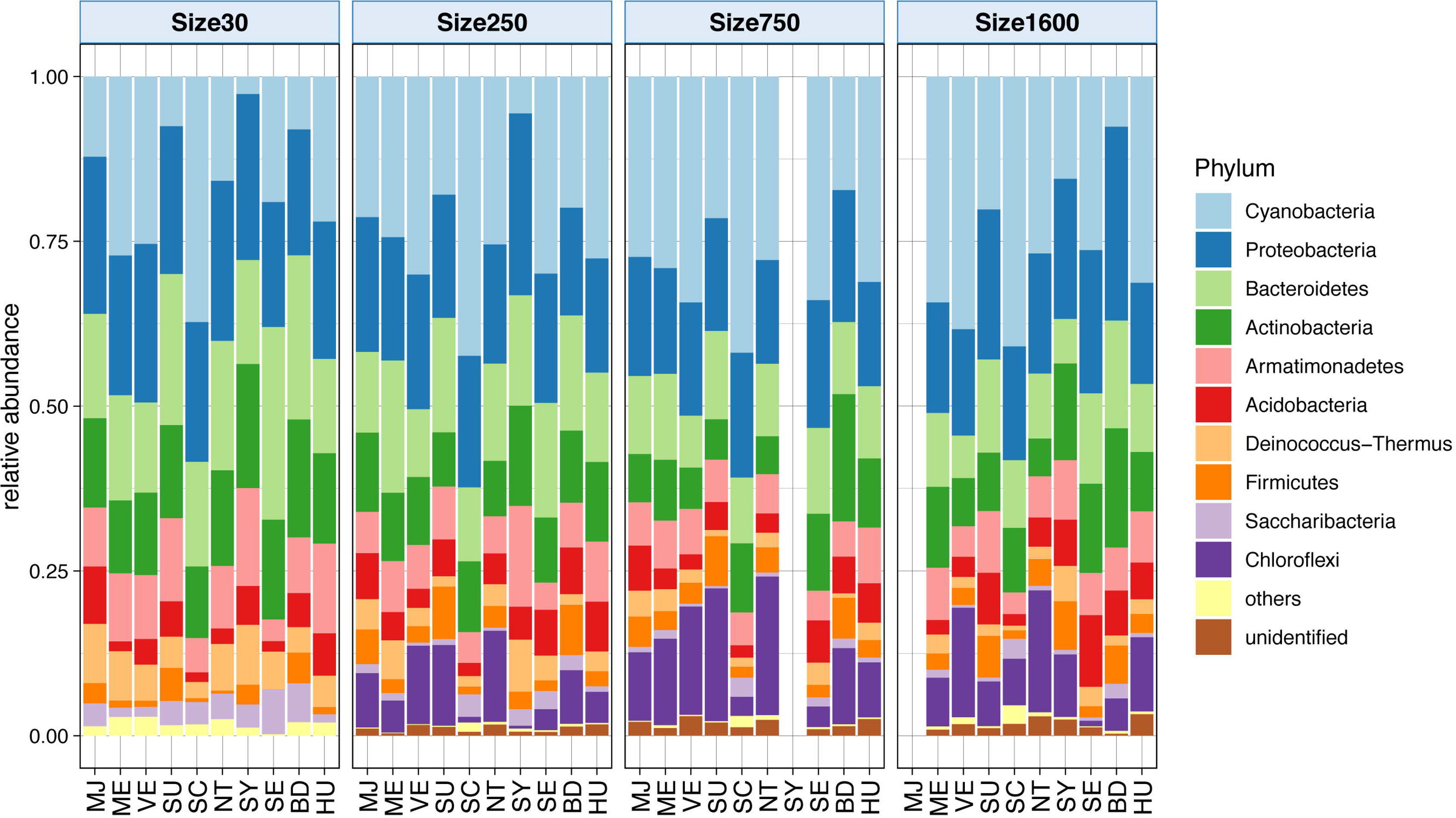
Relative abundance of bacterial phyla of each cryoconite granule size category (Size30, Size250, Size750, Size1600) in 10 glaciers around Qaanaaq.

Homogenously distributed genus across sites and size, which were more than 1% of SIMPER similarities in each size fraction are shown in SI Table 2 and Fig. 2. Genus that constantly high SIMPER similarity across all of the size fractions were *Phormidesmis*, *Acidiphilium*, *Solitalea*, *Granulicella*, *Deinococcus*, *Hymenobacter, Ferruginibacter*. *Salinibacterium* and hgcI_clade in *Sporichthyaceae* tended to be common in smaller size fractions (Size 30-250) and *Polymorphobacter* is ubiquitously high in smallest and largest size fraction. Within these higher SIMPER genus, *Cyanobacteria* genus *Phormidesmis* was most abundant and primary dominated in all size fractions in VE (10-19%) and HU (15-26%), and in the size fractions larger than 250 in other site except SCH (MJ: 17-23%, NT: 17-20%, SER: 18-26%, SUN: 14-15%, ME: 17-18%, SY: 14%, BD: 16%). SIMPER similarity of Phormidesmis among sites largely increase from Size 250 fraction, and other genus such as *Deinococcus*, *Hymenobacter and Ferruginibacter* tend to decrease by size increasing. These homogeneously distributed genus are mainly composed by few ASVs, except *Acidiphilium* which are composed by 5 ASVs (SI Fig. 2). Unique genus at each sites and size fractions, which analyzed by indicator species analysis, are shown in SI Table3. In all size fractions, highest number of indicator genera were detected in SCH and *Leptolyngbya* (Cyanobacteria) is most abundant genus. *Leptolyngbya* and *Aetherobacter* (Proteobacteria) were common in all 4 size fractions of SCH with more than 0.1 frequency of INDVAL (SI Table 3 and Fig. 2). *Leptolyngbya* is mainly composed by three unique ASVs (ASV 10, ASV22 and ASV549) is mostly dominated. In other glacier, *Pedobacter* is only high (more than 0.1) and unique in Size 30 in SER.

**Figure 2:**
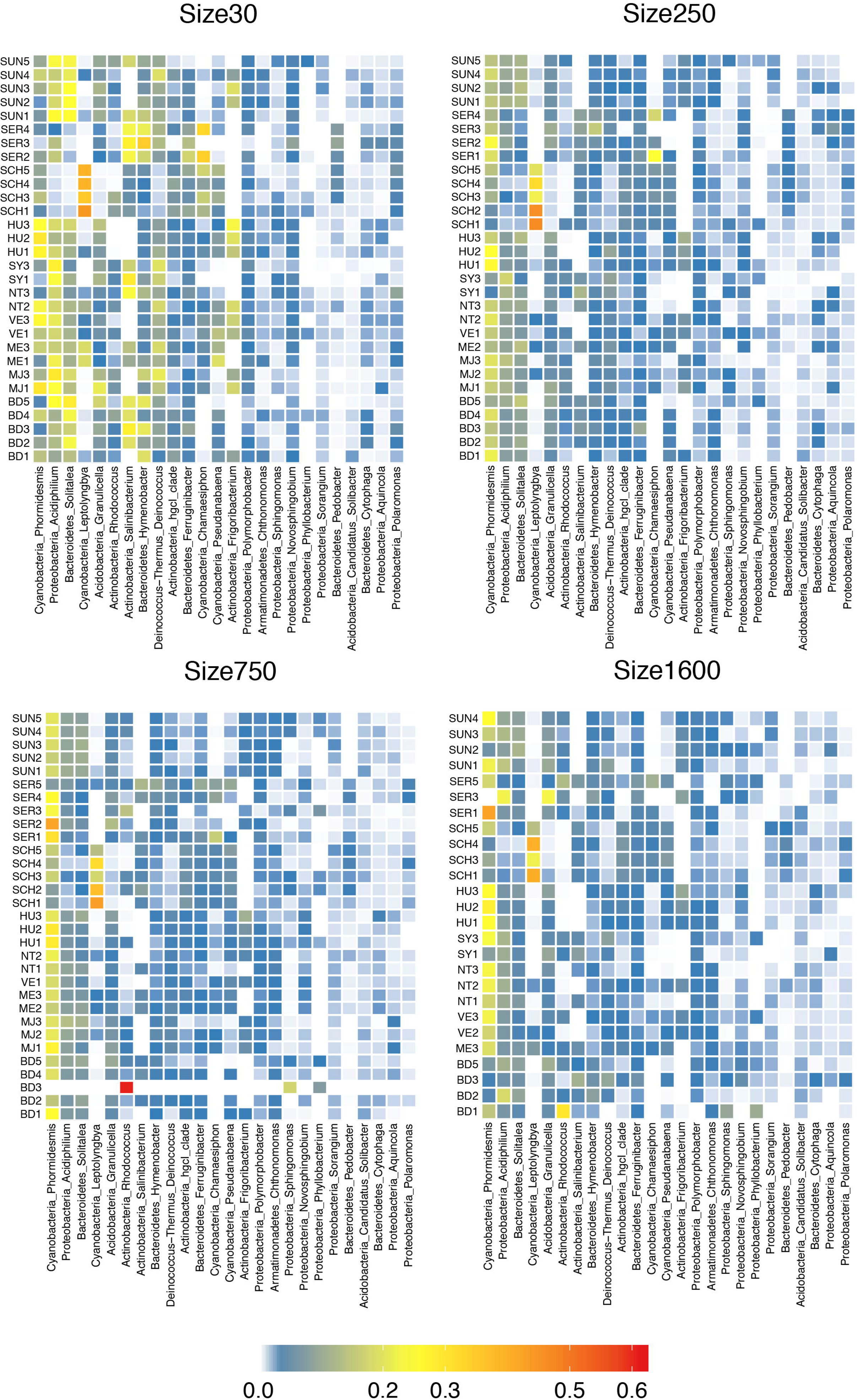
Relative abundance of bacterial genera of each cryoconite granule size category (Size30, Size250, Size750, Size1600) in 10 glaciers around Qaanaaq.

### 2: Diversity

Alpha diversities have strong significant difference among glaciers in Size30 for Chao1 (ANOVA F: 3.99, p<0.005) and in Size 30 for Shannon (ANOVA F: 4.62, p<0.005), and in Size30 and Size 250 for Simpson reciprocal (ANOVA F: 6.66, p<0.0005, 5.88, p<0.0005), and these become less significant or no significance by granule size increasing (Fig. 3). Otherwise, difference of alpha diversities among size in each glacier are not significant by ANOVA. Principal Component Analysis (PCA) of Bray Curtis dissimilarity shows SCH and SER are tend to be different from other glacier in all size fractions (SI Fig. 3). Differences of beta diversities (Bray Curtis and weighted UniFrac) among glaciers were high in small size fraction (ANOSIM_R: 0.818 and 0.679) and gradually decrease to 0.576 and 0.508 in Size 750, then slightly increase to 0.580 and 0.521 in Size 1600 (Fig. 4 a).

**Figure 3:**
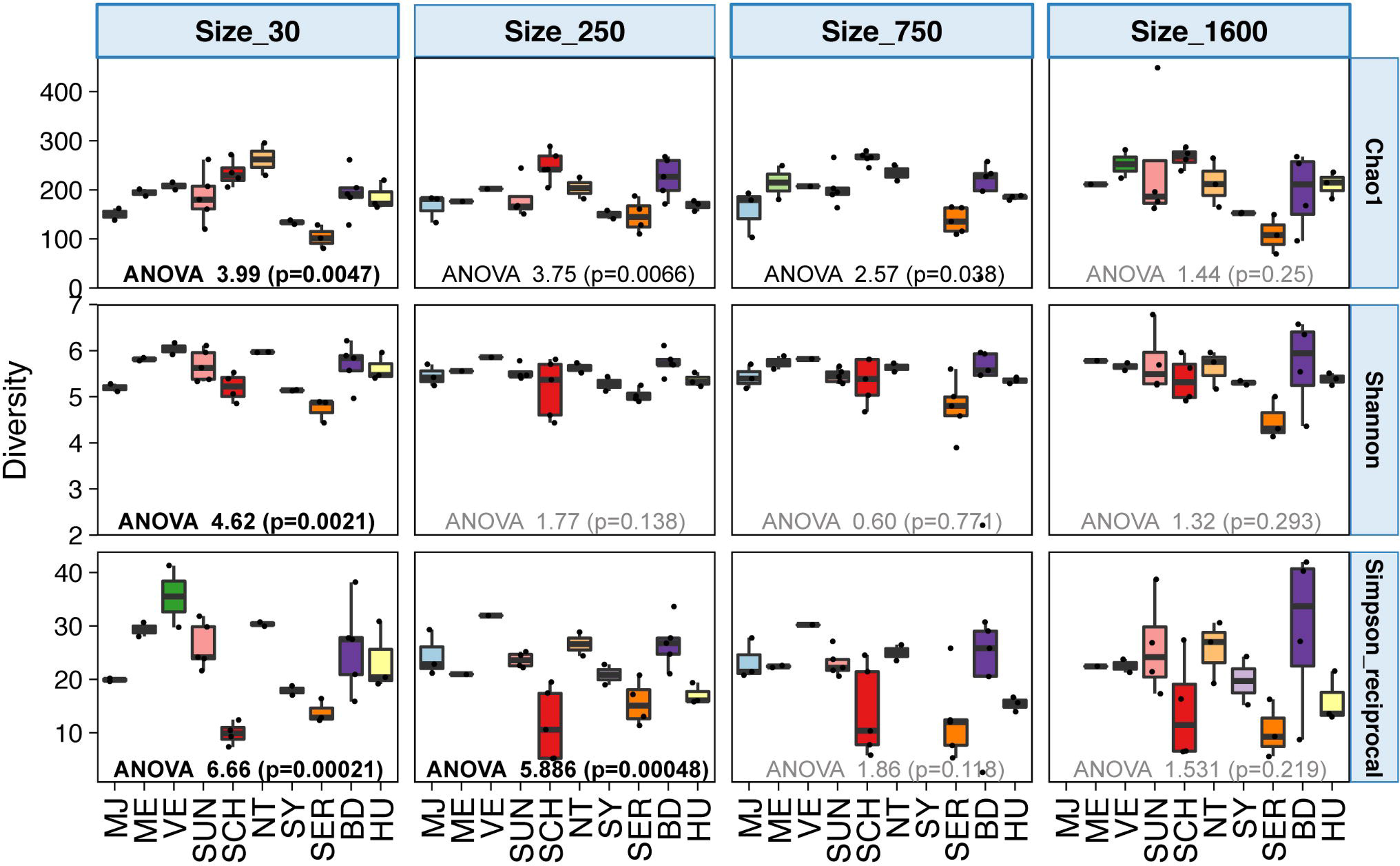
Alpha diversities (Chao1, Shannon and Simpson reciprocal) of each cryoconite granule size category (Size30, Size250, Size750, Size1600) in 10 glaciers around Qaanaaq. Results of ANOVA among all glaciers are shown in each.

**Figure 4:**
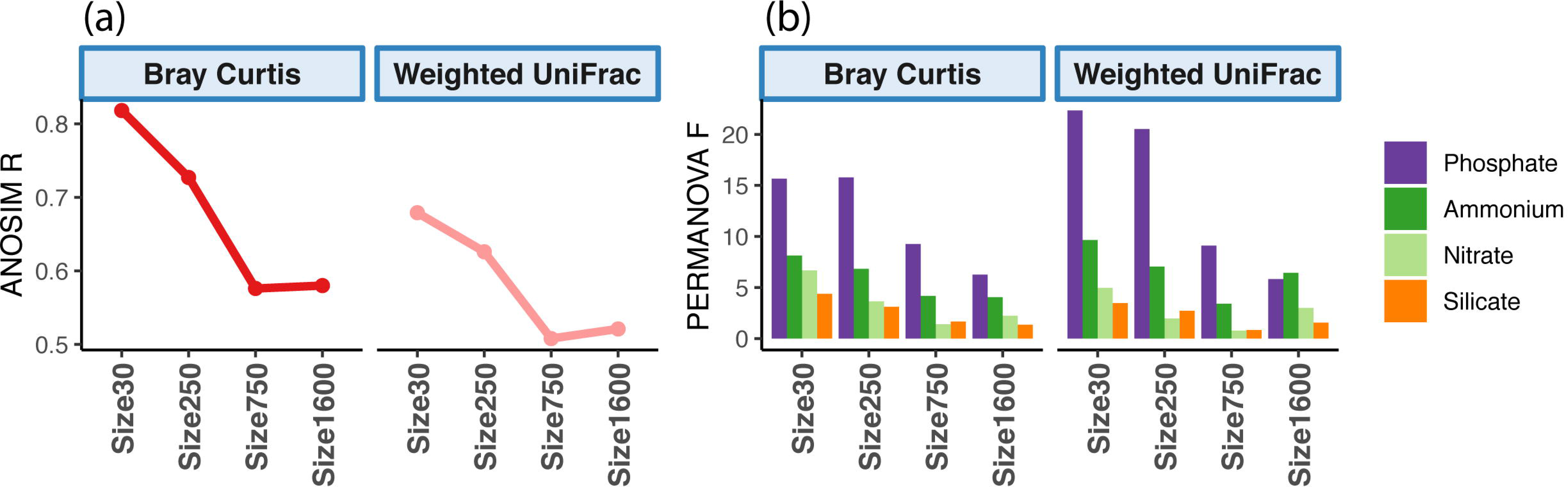
Dissimilarities of bacterial communities among all glaciers based on of Bray Curtis and Weighted UniFrac distances in each granule size category. (a) Dissimilarities was estimated by ANOSIM. (b) Dissimilarities on the effects of nutrients (ammonium, nitrate, phosphate, silicate) was estimated by PERMANOVA.

### 3: Nutrients

#### 3-1: Nutrients in precipitation and meltwater in cryoconite hole

Concentration of nutrients (ammonium, nitrate, phosphate and silicate) in meltwater of CH and precipitation was shown in Fig.5. All nutrients across glaciers (both include/excluding precipitation) were significantly variable (one way ANOVA, p<0.0001). Concentrations of both nitrate and ammonium in precipitation (283-707 ppb, 43-59 ppb) were significantly higher than those in meltwater of CH by Tukey HSD test (p<0.0001). The concentration of silicate in precipitation (17-21 ppb) was significantly lower than meltwater at the sites SY (20-25 ppb) and HU (23-25 ppb), but higher at the other sites. The concentrations of phosphate in SCH (3 - 23 ppb) is significantly higher than other glacier and precipitation (Turkey’s test: p<0.05) except SER (4 - 6 ppb).

**Figure 5:**
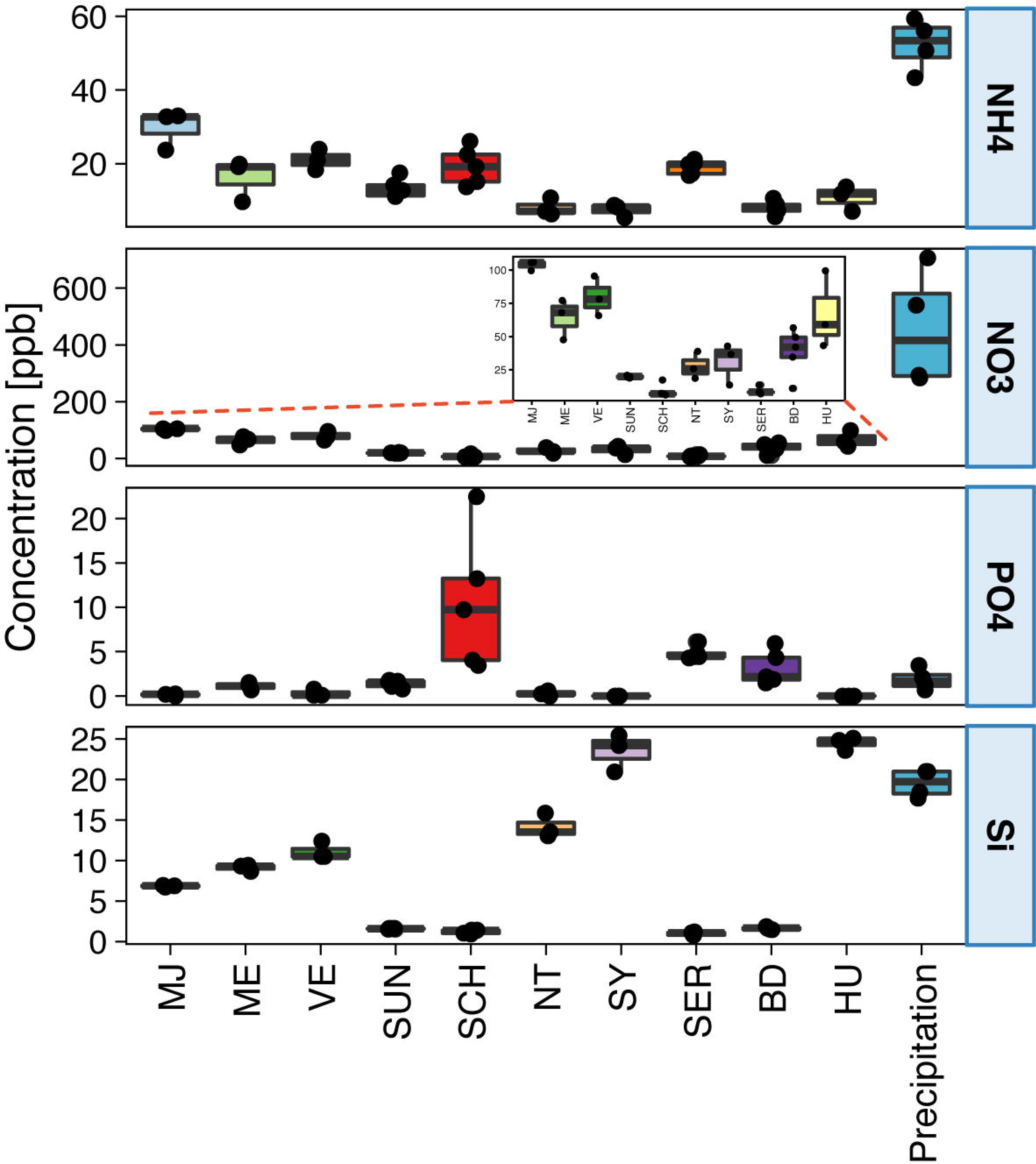
Result of nutrient concentrations (ammonium, nitrate, phosphate, silicate) in 10 glaciers and precipitation sample from Qaanaaq Glacier.

#### 3-2: Correlation between bacterial communities and nutrients in melt water

Multivariate analysis (PERMANOVA) shows that contribution of nutrients concentration on the beta diversity (Bray Curtis and weighted UniFrac) differed among the size fractions of granules (SI Table 4, Fig. 4 b). Phosphate is best factor to explain the dissimilarity of bacterial communities (Pseudo F: 15.7, 22.3, respectively), and Pseudo-F of all nutrients gradually decreased with increase of the granule size. Correlation table between nutrients and genus in Fig. 6 shows *Phormidesmis* and *Frigoribacterium* were positively correlated with nitrate in smaller size fractions. And *Leptolyngbya*, *Pedobacter* and hgcI_clade were positively correlated with phosphate in mostly all size fraction, however, phosphate is negatively correlated with *Acidiphilium*, *Granulicella*, *Deinococcus* in smaller size fractions.

## Discussion

### 1: Cyanobacteria characterizing cryoconite granules

On all of the study glaciers, filamentous cyanobacteria genus *Phormidesmis* (mostly *P. priestleyi*) was abundant in all size fractions of granules. *P. priestleyi* has been isolated from Antarctic lake originally (Taton et al. 2010), and has also been reported from many of Arctic glaciers (Chrismas et al. 2016, Gokul et al. 2016, Uetake et al. 2016, Anesio et al. 2017, Cameron et al. 2017). Relative abundance and SIMPER similarity of this genus largely increased in larger than size 250, which Uetake *et al.* (2016) pointed out as the distinct boundary of carbon ratio and species diversity in cryoconite granules (SI Table 3, Fig. 2). Since cryoconite granules are enlarged by growth of filamentous cyanobacteria (Hodson et al. 2010; Langford et al. 2010), dominated filamentous cyanobacteria *P. priestleyi* is the key species to understand formation process of cryoconite granules on Arctic glaciers (Gokul et al. 2016, Uetake et al. 2016, Anesio et al. 2017). Moreover, major 3 ASVs (ASV1, ASV5 and ASV7 in SI Fig. 2) are corresponded to the cyanobacteria OTU1 in Segawa et al. (2017), which was widely distributed from polar to Asian glaciers, with high identities 99.3％ (399/405), 100% (405/405), 99.8% (404/405) respectively. This phylotype (OTU1) was reported as major cyanobacterium (mostly more than 50% of total cyanobacteria) on glaciers in Svalbard, western parts of Greenland, and Alaska, this result is well corresponding with the high relative abundance of *P.priestleyi* in this study.

Despite of the high abundance of this genus, 16S rRNA gene network analysis showed that abundance of *P. priestleyi* are poorly connected to other bacteria (Gokul et al. 2016). However, even if the correlations between the bacterial species were weak, *P.priestleyi* is primary important as an ecosystem engineer in terms of creating habitable space for other microorganisms by granulation (Gokul et al. 2015, Musilova et al. 2015), and other high SIMPER genus would be general habitants in this system.

Other major filamentous cyanobacteria, which can be the main frame of the structure of cryoconite granules, were *Psudoanabena* and *Leptolyngbya*. The SIMPER similarity for *Psudoanabena* was relatively higher in Size30, Size250 and Size750, however, it decreased as the granule size increased and absent in size 1600. Therefore, it is unlikely to contribute mainly the formation of cryoconite granules. *Leptolyngbya* was a relatively rare genus at all sites except SCH, where the number of indicator species was highest and the beta diversity was very different from other glaciers (SI Table3 and SI Fig.3). The major 3 ASVs belonging to *Leptolyngbya* (ASV10, ASV22 and ASV549 in SI Fig. 2) correspond to the phylotypes of Segawa’s OTU7 (99.9%), OTU0 (100%), OUT7 (99.8%), respectively, which are major in Asian (OTU0) and polar regions (OTU7) in Segawa et al. (2017). Despite of a large geographical separation, this result may suggest relationship among SCH and previous studied glacier (Rink Ice cap in Antarctica Peninsula and Foxfonna Glacier in Svalbard), where Segawa’s OTU7 is most abundant. The factors crucial for generating this similarity are still uncertain, but comparison of the geographical variations of environmental factors and bacterial community structure in worldwide may be helpful to understand.

### 2: Variations in bacterial diversity and susceptibility for nutrients

#### Effect of nutrients on microbial community in different size of cryoconite granules

The change of microbial communities in different granule size was variable even between the outlet glaciers of the same ice cap or ice sheet (Qaanaaq ice cap: SCH, SYand SER, Greenland Ice Sheet: MJ, ME, SUN, NT. BD and HU). Microbial communities became more homogeneous across the glaciers with increasing of granule size. The degree of dissimilarities of bacterial communities across the glaciers estimated by two different method (Bray Curtis and weighted UniFrac) were smaller for larger granule size and variability of alpha diversities tested by one way ANOVA (F-value) are higher in smaller granule (Fig. 4). This suggests that the small size fractions contain more unique ASV than more matured larger size. Correlation coefficients and their significance between major genus and nutrients largely changed in each size fraction, suggesting that nutrients susceptibility to genus changed with size of granule. Because, microbial cells and other substances are loosely tightened in smaller granules, bacterial cells are likely to be affected directly by meltwater than those in larger granules, which microbial cells and/or substances (e.g. extracellar polysachaloides) are densely aggregated. In contrast, in larger granules, nutrients such as ammonium are produced and supplied by nitrogen fixation in anoxic condition inside of granule (Telling et al. 2012), and nitrate would be supplied by nitrification and removed by denitrification processes (Segawa et al. 2014). Althought relationship between bacteria communities and nutrients in CHs was affected by meltwater inflow and outflow in normal year. Supraglacial melt water flow was very limited, because summer in 2013 was much colder than usual (Tedstone et al. 2017) and the ablation ice surface, where cryoconite granules are usually exposed (e.g. middle of Qaanaaq Glacier in Uetake et al. 2016), was still covered with thick snow and/or superimposed ice throughout the ablation season. Therefore, we assumed melt water effect would be less important than that in high melting year and will discuss further more about each nutrient in next paragraphs.

#### Nitrate

Nitrate is one of most influential and well reported nutrients in supraglacial microbial community, because that supports the activity and growth of microorganisms and its input will be expected increasing in the future (Telling et al. 2012, Cameron et al. 2017). In this study, nitrate as well as ammonium are significantly much higher in precipitation, nitrate was assumed as anthropogenic influence (Chellman et al. 2016) and that will be affect bacterial communities on the glaciers. Genera *Phormidesmis* and *Frigoribacterium* were positively correlated with nitrate in Size 30 - 250 and Size 30 - 750, respectively (Fig. 6). Despite of strong anthropogenic nitrate input from deposition, nitrate concentration widely vary among glaciers with significant difference (ANOVA P < 0.0001), and glacier MJ, ME, VE and HU are tended to significantly different form other glaciers (TurkeyHSD). Since relative abundance of *Phormidesmis* in Size 30 is especially high in these higher nitrate concentrated glaciers (Fig. 2), nitrate is likely to promote the growth of *Phormidesmis* in early granulation process. Short term (6 weeks) in situ incubation experiments in Greenland showed that the artificial supply of nitrate onto the cryoconite did not affect the bacterial community (Cameron et al. 2017). On the other hand, long term (1-3 years) microcosms experiment controlled nitrogen and phosphate input showed significant increase of carbon concentration. Difference between these 2 studies may be caused by duration of incubation, because growth rate of bacteria and cryoconite granules are very slow due to cold temperature (e.g. 3.5-7 years need for mature cryoconite granule in Chinese glacier: Takeuchi et al. 2010), and microbial communities would be not be able to utilize nutrients enough for sudden change of supply. Also sequencing data without granule size sorting would be strongly affected by larger granules, which include more biomass and DNA than smaller size, therefore, nutrient and microbe interaction would be masked by majority of microbial communities in large granule.

Contrary to small granule, effect of nitrate was minimum in largest size fraction. That was possibly due to nitrate supply by nitrification from granule bacterial communities (Segawa et al. 2014), however, typical bacteria genera related to denitrification (such as *Nitrosomonas*, *Nitrosospira*, *Nitrosococcus*, *Nitrobacter*, *Nitrospina*, *Nitrococcus*, *Nitrosoglobus*, *Nitrospira*) were not detected, and taxonomy-based function estimate also shows very few abundances of ammonia monooxygenase genes (amoA, amoB and amoC) than nitrate reductase (SI Fig. 4). However, heterogeneous distribution of nitrate, which are homogeneously deposited from air, would be caused by nitrogen cycling driven by microorganisms. Therefore, in order to understand relationship between microorganisms and nitrate, we should test more robust evidence from measurements using oxygen and nitrogen stable isotopes of nitrate (Segawa et al. 2014, Chellman et al. 2016) and gene expression of nitrogen cycle genes and response of *P.priestlreyi* under various incubation conditions.

#### Phosphate

Phosphate is also recognized as another important nutrient on glacier microbial habitat (Musilova *et al.* 2016). This is probably accumulated as non-bioavailable organophosphorus compounds, however, glacier microorganisms are able to degrade these to phosphate by using phosphatase (Stibal *et al.* 2009). Phosphate concentration is significantly high in site SCH, where the bacterial community was distinctive. Genus positively correlated with phosphate is *Leptolyngbya* (Cyanobacteria), *Pedobacter* (Bacteroidetes) and hgcI_clade (Actinobacteria), and all of these are relatively high in SCH. Among these genera, *Leptolyngbya* is most abundant genus in SCH and positively correlated with phosphate in all size fractions. Although 3 ASVs from this genus were closely related to many polar and alpine sequences, there is no information about phosphate requirement for glacier *Leptolyngbya*. However, phosphate metabolism, uptake and transport genes were detected in alpine glacier cryoconite (Edwards *et al.* 2013), and this study indicate phosphate cycling by microorganisms are abundant in glacier microbial ecosystem. Among these phosphate genes, genes related to phosphate sensor and response (PhoR, PhoB: Santos-Beneit, 2015) were tended to higher in phosphate high glacier such as SCH, SER and BD (SI Fig. 5), and this may also support phosphate influence to microbial communities.

**Figure 6:**
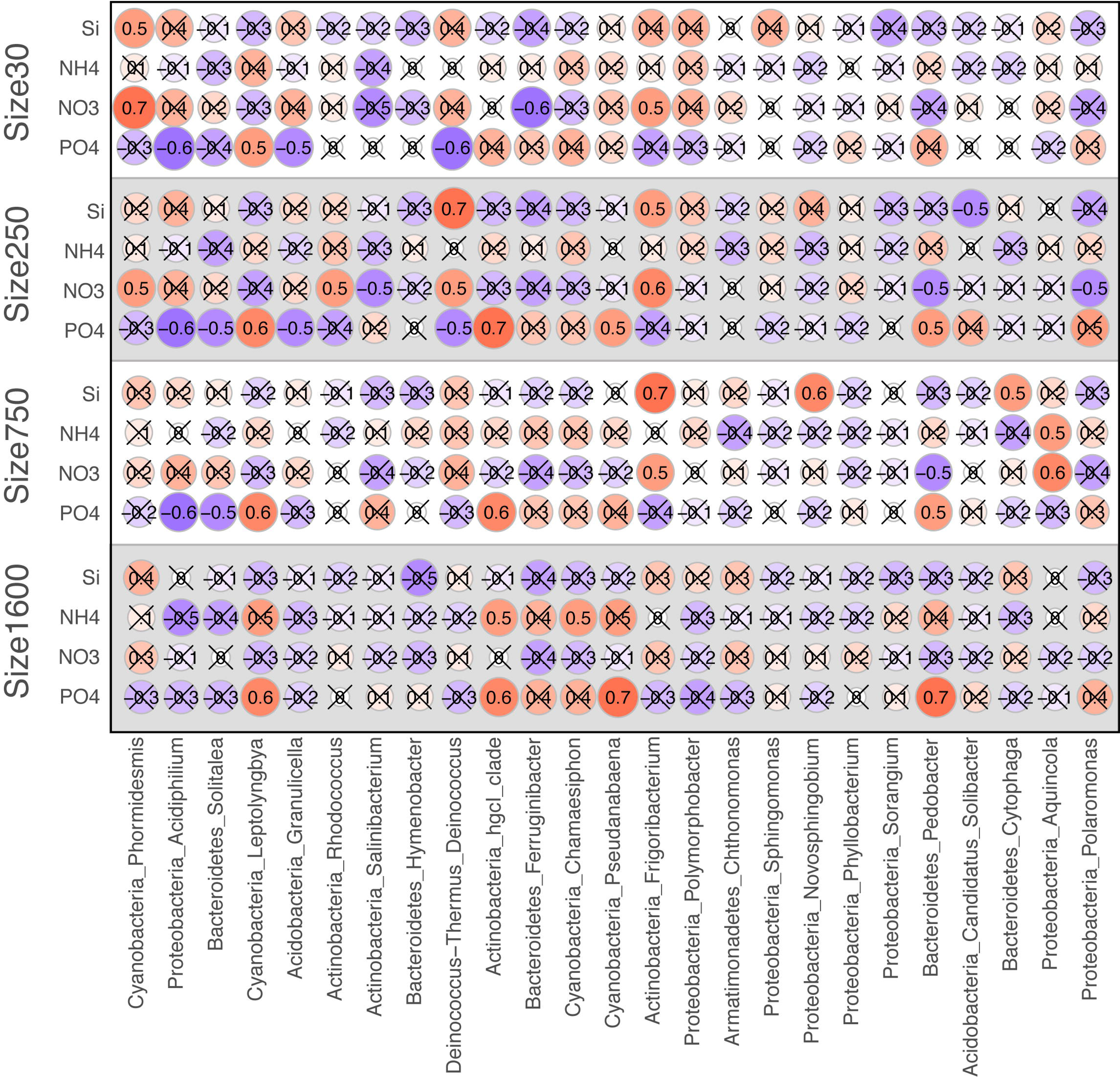
Correlation between 50 major genera and nutrients in each size category of granule. Correlation coefficient without significance (P > 0.01) was crossed out.

Although relationship between glacier microorganisms and phosphate was not well understood yet, recent study showed phosphate was much stronger factor to change microbial communities than nitrogen in early succession stage of glacier foreland (Darcy *et al.* 2018). Therefore, phosphate is also important factor for glacier microbial ecosystem showing in this study. However, mechanisms causing significant difference of phosphate concentration among glaciers is still question. Since the input of phosphate to the glacier surface by precipitation is obviously low (SI Table 4, Fig. 1), and origin of phosphate is from minerals on each glacier, because microorganisms contribute to dissolution of insoluble secondary phosphates in mineral (Banfield *et al.* 1999), thus the mineral composition may affect phosphate abundance on glacier. Phosphate elution process on the glacier surface is also not understood well, but phosphate is high in turbid subglacial meltwater (Hodson *et al.* 2004, Mindl *et al.* 2007). Relationship between bacteria communities and minerals has been reported in previous studies, which show the interactions between microbial community and mineral particles on glacier in Svalbard (Edward *et al.* 2011). Also relationship between the abundance of cryoconite and outcropped minerals (Nagatsuka *et al.* 2016) was reported in Uetake *et al.* 2016. Composition of minerals and their solute have potential to affect the microbial growth in some special site (SCH in this study), however, more measurements such as elemental analysis using ICP-MS (Lutz *et al.* 2014, 2016) and elution potential of nutrients by laboratory experiments are needed in the future study.

## Acknowledgement

The authors thank to member of “Snow Impurity and Glacial Microbe effects on abrupt warming in the Arctic (SIGMA)” project and “Green Network of Excellence (GRENE)” program in 2013, especially Dr. Akane Tsushima in Research Institute for Humanity and Nature, and Dr. Shun Tsutaki in University of Tokyo for field sampling. Special thanks are due to Mr. Tetsuhide Yamasaki and Mr. Finn Hansen for general support of fieldwork and Ms. Sakiko Daorana for logistics and accommodation in Qaanaaq village. We also thank Mr. Kenichi Watanabe and Ms. Mizuho Mori for assistance in laboratory experiments and Dr. Justin Lee in Colorado State University for English expressions.

Conflict of interest. None declared.

