## Supplemental Table 1 for "Bacterial community changes with cryoconite granule size and their susceptibility to exogenous nutrients on 10 glaciers in northwestern Greenland"

| **SI Table 1** | | |  |  | |
| --- | --- | --- | --- | --- | --- |
| **Information about sampling sites on 10 glaciers.** | | |  |  | |
| Glacier | Abbreviation | Coordinate | | | Altitude (m) |
| Morris Jesup Glacier | MJ | 77°53′16″ N, 71°08′21″ W | | | NA |
| Meehan Glacier | ME | 77°52′28″ N, 70°18′46″ W | | | NA |
| Verhoeff Glacier | VE | 77°51′38″ N, 69°52′08″ W | | | NA |
| Sun Glacier | SUN | 77°46′36″ N, 69°27′18″ W | | | 102 |
| Scarlet Heart Glacier | SCH | 77°39′56″ N, 69°27′03″ W | | | 140 |
| No name glacier next of Tugoto Glacier | NT | 77°43′29″ N, 69°02′48″ W | | | NA |
| Syd Glacier | SY | 77°33′37″ N, 68°42′17″ W | | | NA |
| Sermiarssupaluk | SER | 77°30′05″ N, 68°45′24″ W | | | NA |
| Bowdoin Glacier | BD | 77°40′09″ N, 68°34′58″ W | | | 48 |
| Hubbard Glacier | HU | 77°32′17″ N, 67°49′19″ W | | | NA |
