## Supplemental Table 2 for "Bacterial community changes with cryoconite granule size and their susceptibility to exogenous nutrients on 10 glaciers in northwestern Greenland"

| **SI Table2** | | | |  | | |
| --- | --- | --- | --- | --- | --- | --- |
| **List of genera with higher similarity percentage (SIMPER > 1.0) in three categories (1: All size is for genera have higher similarity throughout all sizes, 2: Small size is for genera have higher similarity only in smaller granules, and 3: others is for genera was excluded with above 2 categories)** | | | | | | |
| Category | taxonomy | Size | Av.Abund | | Av.Sim | Contrib% |
| All size | Bacteria;Cyanobacteria;Cyanobacteria;SubsectionIII;FamilyI;Phormidesmis | 30 | 0.06 | | 3.79 | 7.33 |
|  |  | 250 | 0.15 | | 11.6 | 21.22 |
|  |  | 750 | 0.19 | | 14.73 | 27.5 |
|  |  | 1600 | 0.16 | | 10.46 | 21.93 |
|  | Bacteria;Proteobacteria;Alphaproteobacteria;Rhodospirillales;Acetobacteraceae;Acidiphilium | 30 | 0.08 | | 5.4 | 10.44 |
|  |  | 250 | 0.08 | | 5.3 | 9.69 |
|  |  | 750 | 0.06 | | 4.35 | 8.11 |
|  |  | 1600 | 0.08 | | 4.81 | 10.08 |
|  | Bacteria;Bacteroidetes;Sphingobacteriia;Sphingobacteriales;Sphingobacteriaceae;Solitalea | 30 | 0.07 | | 3.9 | 7.55 |
|  |  | 250 | 0.07 | | 4.11 | 7.53 |
|  |  | 750 | 0.05 | | 3.24 | 6.05 |
|  |  | 1600 | 0.05 | | 2.95 | 6.18 |
|  | Bacteria;Acidobacteria;Acidobacteria;Acidobacteriales;Acidobacteriaceae_(Subgroup_1);Granulicella | 30 | 0.04 | | 2.23 | 4.32 |
|  |  | 250 | 0.05 | | 3.24 | 5.94 |
|  |  | 750 | 0.04 | | 2.28 | 4.26 |
|  |  | 1600 | 0.05 | | 2.68 | 5.62 |
|  | Bacteria;Deinococcus-Thermus;Deinococci;Deinococcales;Deinococcaceae;Deinococcus | 30 | 0.05 | | 3.53 | 6.82 |
|  |  | 250 | 0.03 | | 1.68 | 3.07 |
|  |  | 750 | 0.02 | | 1.07 | 2 |
|  |  | 1600 | 0.02 | | 1.11 | 2.32 |
|  | Bacteria;Bacteroidetes;Cytophagia;Cytophagales;Cytophagaceae;Hymenobacter | 30 | 0.05 | | 2.76 | 5.34 |
|  |  | 250 | 0.03 | | 1.53 | 2.8 |
|  |  | 750 | 0.02 | | 1.12 | 2.1 |
|  |  | 1600 | 0.02 | | 1.1 | 2.3 |
|  | Bacteria;Bacteroidetes;Sphingobacteriia;Sphingobacteriales;Chitinophagaceae;Ferruginibacter | 30 | 0.03 | | 1.81 | 3.5 |
|  |  | 250 | 0.02 | | 1.45 | 2.65 |
|  |  | 1600 | 0.02 | | 1.34 | 2.8 |
| Small size | Bacteria;Actinobacteria;Actinobacteria;Frankiales;Sporichthyaceae;hgcI_clade | 30 | 0.02 | | 1.29 | 2.5 |
|  | Bacteria;Actinobacteria;Actinobacteria;Micrococcales;Microbacteriaceae;Salinibacterium | 30 | 0.06 | | 2.31 | 4.47 |
|  |  | 250 | 0.03 | | 1.37 | 2.51 |
| Other | Bacteria;Proteobacteria;Alphaproteobacteria;Sphingomonadales;Sphingomonadaceae;Polymorphobacter | 30 | 0.02 | | 1.16 | 2.24 |
|  |  | 1600 | 0.02 | | 1.12 | 2.34 |
