## Supplemental Table 3 for "Bacterial community changes with cryoconite granule size and their susceptibility to exogenous nutrients on 10 glaciers in northwestern Greenland"

| **SI Table3** | | | |  | |
| --- | --- | --- | --- | --- | --- |
| **List of genera, which are estimated as indicator genera for each granule size using indicator species analysis (indval function of “labdsv” packages of R).** | | | |  | |
| Size30 | group | indval | p-value | | frequency |
| Bacteria.Bacteroidetes.Cytophagia.Cytophagales.Cyclobacteriaceae.NA | NT | 0.8529103 | 0.007 | | 0.005 |
| Bacteria.Proteobacteria.Alphaproteobacteria.Rhizobiales.Bradyrhizobiaceae.Rhodopseudomonas | HU | 0.6086561 | 0.005 | | 0.020 |
| Bacteria.Proteobacteria.Alphaproteobacteria.Rhodospirillales.Rhodospirillaceae.Inquilinus | HU | 0.561228 | 0.005 | | 0.006 |
| **Bacteria.Cyanobacteria.Cyanobacteria.SubsectionIII.FamilyI.Leptolyngbya** | **SCH** | **0.5592874** | **0.004** | | **4.827** |
| **Bacteria.Proteobacteria.Deltaproteobacteria.Myxococcales.Polyangiaceae.Aetherobacter** | **SCH** | **0.9800778** | **0.001** | | **0.271** |
| **Bacteria.Bacteroidetes.Sphingobacteriia.Sphingobacteriales.Chitinophagaceae.Parasediminibacterium** | **SCH** | **0.8953339** | **0.001** | | **0.184** |
| **Bacteria.Cyanobacteria.Cyanobacteria.SubsectionIII.FamilyI.Tychonema** | **SCH** | **0.8737301** | **0.001** | | **0.145** |
| Bacteria.Gemmatimonadetes.Gemmatimonadetes.Gemmatimonadales.Gemmatimonadaceae.Gemmatimonas | SCH | 0.9040302 | 0.001 | | 0.082 |
| Bacteria.Proteobacteria.Betaproteobacteria.Burkholderiales.Comamonadaceae.NA | SCH | 0.5728536 | 0.002 | | 0.081 |
| Bacteria.Verrucomicrobia.Opitutae.Opitutales.Opitutaceae.Opitutus | SCH | 0.6490304 | 0.001 | | 0.034 |
| Bacteria.Armatimonadetes.Fimbriimonadia.Fimbriimonadales.Fimbriimonadaceae.NA | SCH | 0.9397069 | 0.001 | | 0.022 |
| Bacteria.Bacteroidetes.Cytophagia.Cytophagales.Cytophagaceae.NA | SCH | 0.8831973 | 0.005 | | 0.011 |
| Bacteria.Parcubacteria.Candidatus_Campbellbacteria.NA.NA.NA | SCH | 0.8715931 | 0.001 | | 0.010 |
| Bacteria.Planctomycetes.vadinHA49.NA.NA.NA | SCH | 0.9384729 | 0.001 | | 0.008 |
| **Bacteria.Bacteroidetes.Sphingobacteriia.Sphingobacteriales.Sphingobacteriaceae.Pedobacter** | **SER** | **0.5207441** | **0.003** | | **0.657** |
| Bacteria.Actinobacteria.Actinobacteria.Frankiales.Nakamurellaceae.Nakamurella | SUN | 0.4023288 | 0.001 | | 0.043 |
| Bacteria.Firmicutes.Clostridia.Clostridiales.Clostridiaceae_1.Clostridium_sensu_stricto_13 | SUN | 0.7152954 | 0.003 | | 0.035 |
| Size250 | group | indval | p-value | | frequency |
| Bacteria.Bacteroidetes.Cytophagia.Cytophagales.Cyclobacteriaceae.NA | NT | 0.8222538 | 0.007 | | 0.002 |
| **Bacteria.Cyanobacteria.Cyanobacteria.SubsectionIII.FamilyI.Leptolyngbya** | **SCH** | **0.7692661** | **0.001** | | **4.998** |
| **Bacteria.Proteobacteria.Deltaproteobacteria.Myxococcales.Polyangiaceae.Aetherobacter** | **SCH** | **0.9966102** | **0.001** | | **0.285** |
| **Bacteria.Gemmatimonadetes.Gemmatimonadetes.Gemmatimonadales.Gemmatimonadaceae.Gemmatimonas** | **SCH** | **0.7755126** | **0.004** | | **0.119** |
| Bacteria.Bacteroidetes.Sphingobacteriia.Sphingobacteriales.Chitinophagaceae.Parasediminibacterium | SCH | 0.9523211 | 0.001 | | 0.079 |
| Bacteria.Cyanobacteria.Cyanobacteria.SubsectionIII.FamilyI.Tychonema | SCH | 0.8175726 | 0.001 | | 0.057 |
| Bacteria.Armatimonadetes.Fimbriimonadia.Fimbriimonadales.Fimbriimonadaceae.NA | SCH | 0.7500667 | 0.003 | | 0.039 |
| Bacteria.Verrucomicrobia.Opitutae.Opitutales.Opitutaceae.Opitutus | SCH | 0.8017718 | 0.001 | | 0.032 |
| Bacteria.Chlamydiae.Chlamydiae.Chlamydiales.Parachlamydiaceae.Candidatus_Protochlamydia | SCH | 0.6134079 | 0.001 | | 0.019 |
| Bacteria.Cyanobacteria.Melainabacteria.Vampirovibrionales.NA.NA | SCH | 0.790197 | 0.001 | | 0.012 |
| Bacteria.Planctomycetes.vadinHA49.NA.NA.NA | SCH | 0.8482507 | 0.001 | | 0.010 |
| Bacteria.Planctomycetes.Planctomycetacia.Planctomycetales.Planctomycetaceae.NA | SCH | 0.7772463 | 0.001 | | 0.009 |
| Bacteria.Proteobacteria.Gammaproteobacteria.Cellvibrionales.Cellvibrionaceae.Cellvibrio | SCH | 0.8 | 0.005 | | 0.005 |
| Size750 | group | indval | p-value | | frequency |
| **Bacteria.Cyanobacteria.Cyanobacteria.SubsectionIII.FamilyI.Leptolyngbya** | **SCH** | **0.8176534** | **0.001** | | **4.220** |
| **Bacteria.Proteobacteria.Deltaproteobacteria.Myxococcales.Polyangiaceae.Aetherobacter** | **SCH** | **0.967726** | **0.001** | | **0.218** |
| **Bacteria.Gemmatimonadetes.Gemmatimonadetes.Gemmatimonadales.Gemmatimonadaceae.Gemmatimonas** | **SCH** | **0.8662629** | **0.002** | | **0.167** |
| Bacteria.Bacteroidetes.Sphingobacteriia.Sphingobacteriales.Chitinophagaceae.Parasediminibacterium | SCH | 0.9393794 | 0.001 | | 0.051 |
| Bacteria.Armatimonadetes.Fimbriimonadia.Fimbriimonadales.Fimbriimonadaceae.NA | SCH | 0.87949 | 0.002 | | 0.041 |
| Bacteria.Verrucomicrobia.Opitutae.Opitutales.Opitutaceae.Opitutus | SCH | 0.7559299 | 0.001 | | 0.035 |
| Bacteria.Microgenomates.Candidatus_Pacebacteria.NA.NA.NA | SCH | 0.85363 | 0.004 | | 0.035 |
| Bacteria.Cyanobacteria.Cyanobacteria.SubsectionIII.FamilyI.Tychonema | SCH | 0.7893924 | 0.001 | | 0.031 |
| Bacteria.Proteobacteria.Alphaproteobacteria.Rhodobacterales.Rhodobacteraceae.Rhodobacter | SCH | 0.8001425 | 0.001 | | 0.016 |
| Bacteria.Cyanobacteria.Melainabacteria.Vampirovibrionales.NA.NA | SCH | 0.9109067 | 0.001 | | 0.014 |
| Bacteria.Parcubacteria.Candidatus_Campbellbacteria.NA.NA.NA | SCH | 0.9564093 | 0.001 | | 0.004 |
| Size1600 | group | indval | p-value | | frequency |
| Bacteria.Planctomycetes.Phycisphaerae.Tepidisphaerales.Tepidisphaeraceae.NA | VE | 0.6876267 | 0.001 | | 0.056 |
| Bacteria.Cyanobacteria.Cyanobacteria.NA.NA.NA | VE | 0.6957373 | 0.007 | | 0.045 |
| Bacteria.Acidobacteria.Acidobacteria.Acidobacteriales.Acidobacteriaceae_.Subgroup_1..NA | VE | 1 | 0.009 | | 0.001 |
| **Bacteria.Cyanobacteria.Cyanobacteria.SubsectionIII.FamilyI.Leptolyngbya** | **SCH** | **0.7521574** | **0.004** | | **4.584** |
| **Bacteria.Bacteroidetes.Sphingobacteriia.Sphingobacteriales.Sphingobacteriaceae.Pedobacter** | **SCH** | **0.6375094** | **0.003** | | **0.386** |
| **Bacteria.Proteobacteria.Deltaproteobacteria.Myxococcales.Polyangiaceae.Aetherobacter** | **SCH** | **1** | **0.001** | | **0.241** |
| **Bacteria.Gemmatimonadetes.Gemmatimonadetes.Gemmatimonadales.Gemmatimonadaceae.Gemmatimonas** | **SCH** | **0.868855** | **0.006** | | **0.234** |
| Bacteria.Microgenomates.Candidatus_Pacebacteria.NA.NA.NA | SCH | 0.9825696 | 0.001 | | 0.085 |
| Bacteria.Bacteroidetes.Sphingobacteriia.Sphingobacteriales.Chitinophagaceae.Parasediminibacterium | SCH | 0.9478741 | 0.001 | | 0.076 |
| Bacteria.Armatimonadetes.Fimbriimonadia.Fimbriimonadales.Fimbriimonadaceae.NA | SCH | 0.7922653 | 0.002 | | 0.067 |
| Bacteria.Verrucomicrobia.Opitutae.Opitutales.Opitutaceae.Opitutus | SCH | 0.8517036 | 0.001 | | 0.055 |
| Bacteria.Chlamydiae.Chlamydiae.Chlamydiales.Parachlamydiaceae.Candidatus_Protochlamydia | SCH | 0.8273836 | 0.002 | | 0.042 |
| Bacteria.Proteobacteria.Alphaproteobacteria.Rhodobacterales.Rhodobacteraceae.Rhodobacter | SCH | 0.7524126 | 0.002 | | 0.021 |
| Bacteria.Cyanobacteria.Melainabacteria.Vampirovibrionales.NA.NA | SCH | 1 | 0.001 | | 0.015 |
| Bacteria.Parcubacteria.Candidatus_Campbellbacteria.NA.NA.NA | SCH | 1 | 0.001 | | 0.015 |
