## Supplemental Table 4 for "Bacterial community changes with cryoconite granule size and their susceptibility to exogenous nutrients on 10 glaciers in northwestern Greenland"

| **SI Table 4** | | | | | | |
| --- | --- | --- | --- | --- | --- | --- |
| **Results of PERMANOVA tests on the effects of nutrients (ammonium, nitrate, phosphate, silicate) on the Bray Curtis and weighted UniFrac distances.** | | | | | | |
|  | Source | df | SS | MS | Pseudo-F | P(perm) |
| Bray_Size30 | PO_4_^3-^ | 1 | 0.95156 | 0.95156 | 15.654 | 0.001 |
|  | NO_3_^-^ | 1 | 0.40515 | 0.40515 | 6.6651 | 0.001 |
|  | Si^4+^ | 1 | 0.26633 | 0.26633 | 4.3813 | 0.001 |
|  | NH_4_^+^ | 1 | 0.49419 | 0.49419 | 8.1299 | 0.001 |
| Bray_Size250 | PO_4_^3-^ | 1 | 1.0499 | 1.0499 | 15.79 | 0.001 |
|  | NO_3_^-^ | 1 | 0.24257 | 0.24257 | 3.6481 | 0.007 |
|  | Si^4+^ | 1 | 0.20694 | 0.20694 | 3.1122 | 0.009 |
|  | NH_4_^+^ | 1 | 0.45409 | 0.45409 | 6.829 | 0.001 |
| Bray_Size750 | PO_4_^3-^ | 1 | 0.90951 | 0.90951 | 9.2618 | 0.011 |
|  | NO_3_^-^ | 1 | 0.13757 | 0.13757 | 1.4009 | 0.217 |
|  | Si^4+^ | 1 | 0.16399 | 0.16399 | 1.67 | 0.132 |
|  | NH_4_^+^ | 1 | 0.41031 | 0.41031 | 4.1783 | 0.004 |
| Bray_Size1600 | PO_4_^3-^ | 1 | 0.81461 | 0.81461 | 6.2655 | 0.001 |
|  | NO_3_^-^ | 1 | 0.2889 | 0.2889 | 2.222 | 0.033 |
|  | Si^4+^ | 1 | 0.17503 | 0.17503 | 1.3462 | 0.213 |
|  | NH_4_^+^ | 1 | 0.52717 | 0.52717 | 4.0547 | 0.001 |
| u_UniFrac_Size30 | PO_4_^3-^ | 1 | 0.52271 | 0.52271 | 22.336 | 0.001 |
|  | NO_3_^-^ | 1 | 0.11592 | 0.11592 | 4.9532 | 0.002 |
|  | Si^4+^ | 1 | 0.081331 | 0.081331 | 3.4753 | 0.012 |
|  | NH_4_^+^ | 1 | 0.22574 | 0.22574 | 9.6459 | 0.001 |
| u_UniFrac_Size250 | PO_4_^3-^ | 1 | 0.60565 | 0.60565 | 20.517 | 0.001 |
|  | NO_3_^-^ | 1 | 0.05801 | 0.05801 | 1.9652 | 0.12 |
|  | Si^4+^ | 1 | 0.080132 | 0.080132 | 2.7146 | 0.042 |
|  | NH_4_^+^ | 1 | 0.20814 | 0.20814 | 7.0509 | 0.001 |
| u_UniFrac_Size750 | PO_4_^3-^ | 1 | 0.48172 | 0.48172 | 9.0993 | 0.015 |
|  | NO_3_^-^ | 1 | 0.040407 | 0.040407 | 0.76326 | 0.522 |
|  | Si^4+^ | 1 | 0.043998 | 0.043998 | 0.83109 | 0.443 |
|  | NH_4_^+^ | 1 | 0.18024 | 0.18024 | 3.4045 | 0.025 |
| u_UniFrac_Size1600 | PO_4_^3-^ | 1 | 0.41056 | 0.41056 | 5.8201 | 0.004 |
|  | NO_3_^-^ | 1 | 0.21158 | 0.21158 | 2.9994 | 0.027 |
|  | Si^4+^ | 1 | 0.10933 | 0.10933 | 1.5499 | 0.171 |
|  | NH_4_^+^ | 1 | 0.4544 | 0.4544 | 6.4416 | 0.001 |
