## Supplementary figures and images for "Bacterial community changes with cryoconite granule size and their susceptibility to exogenous nutrients on 10 glaciers in northwestern Greenland"

### Supplemental Figure 1

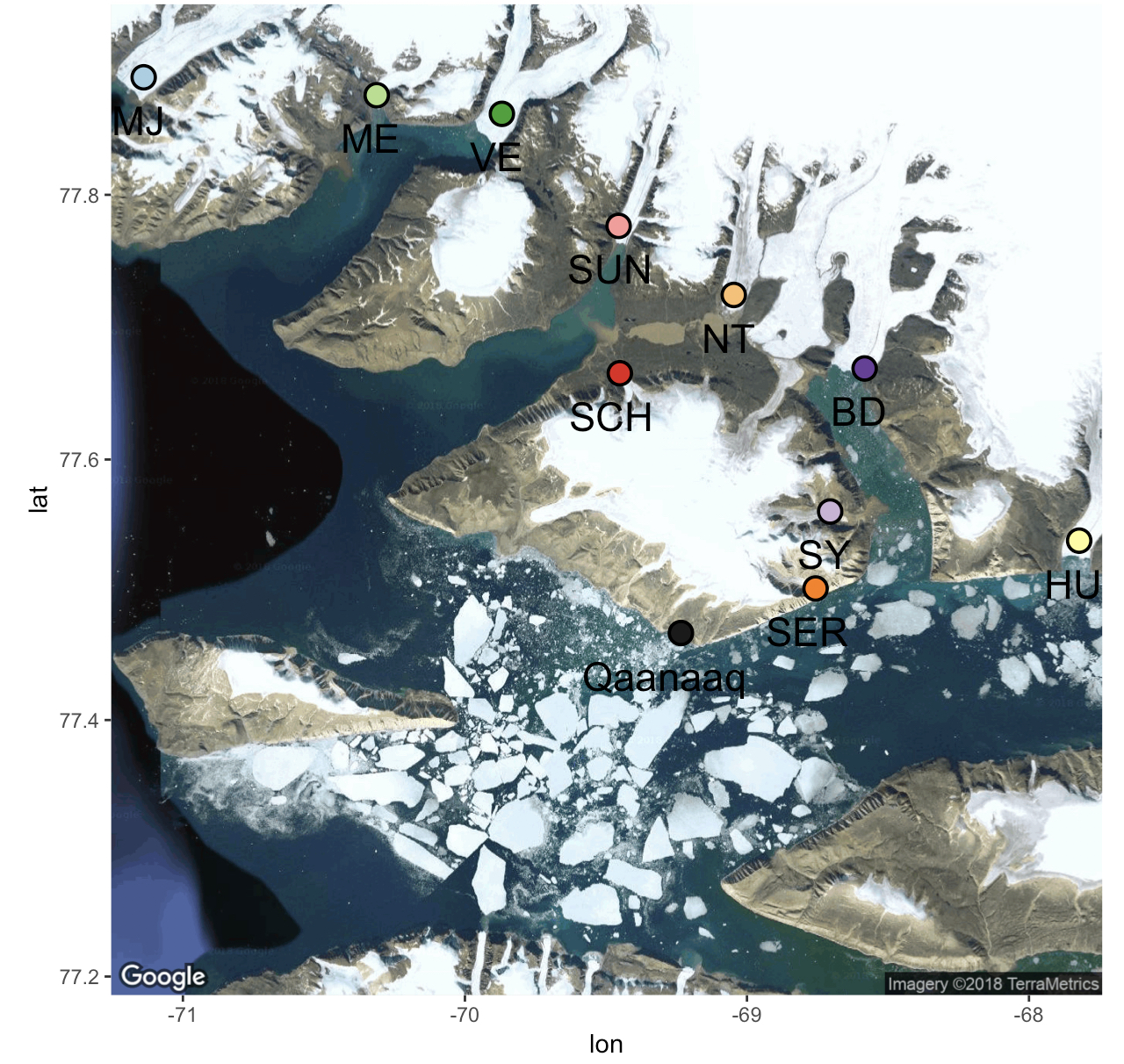

### Supplemental Figure 2

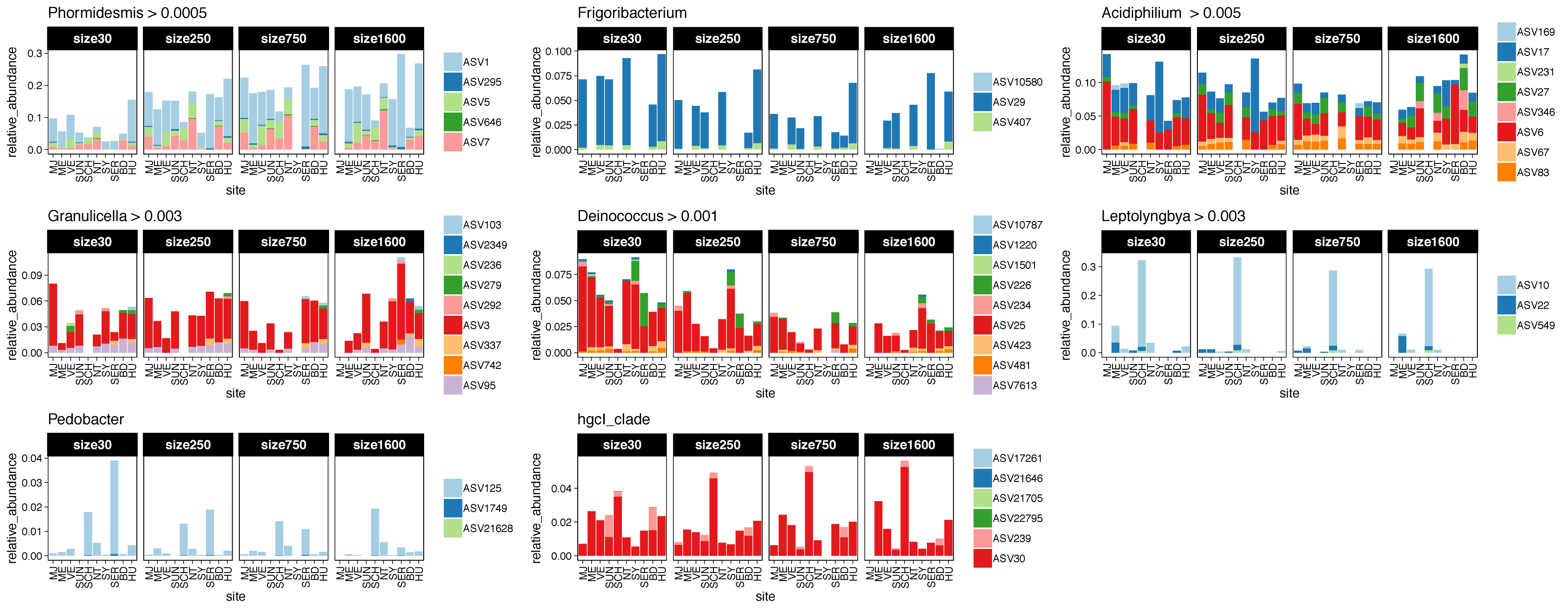

### Supplemental Figure 3

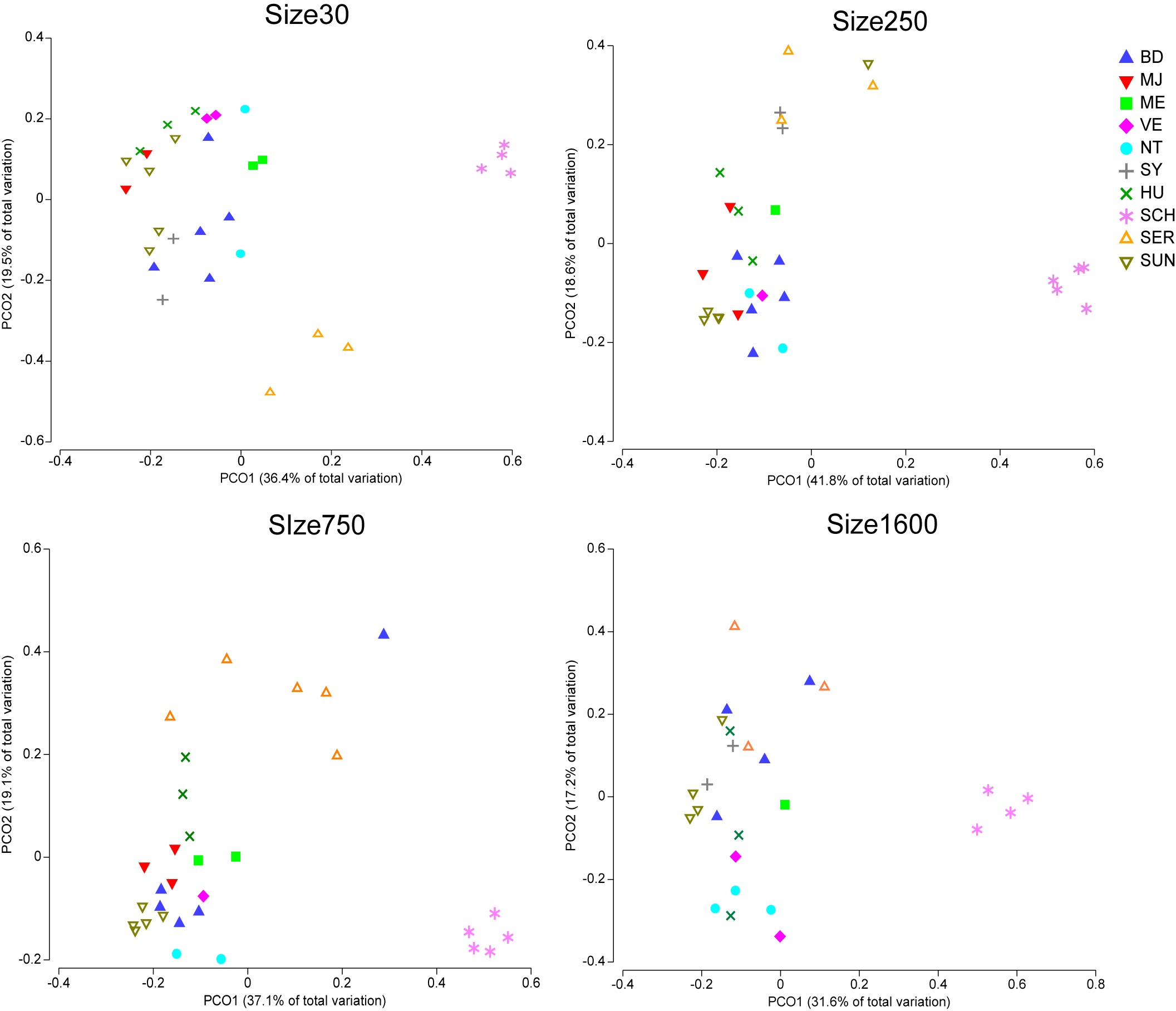

### Supplemental Figure 4

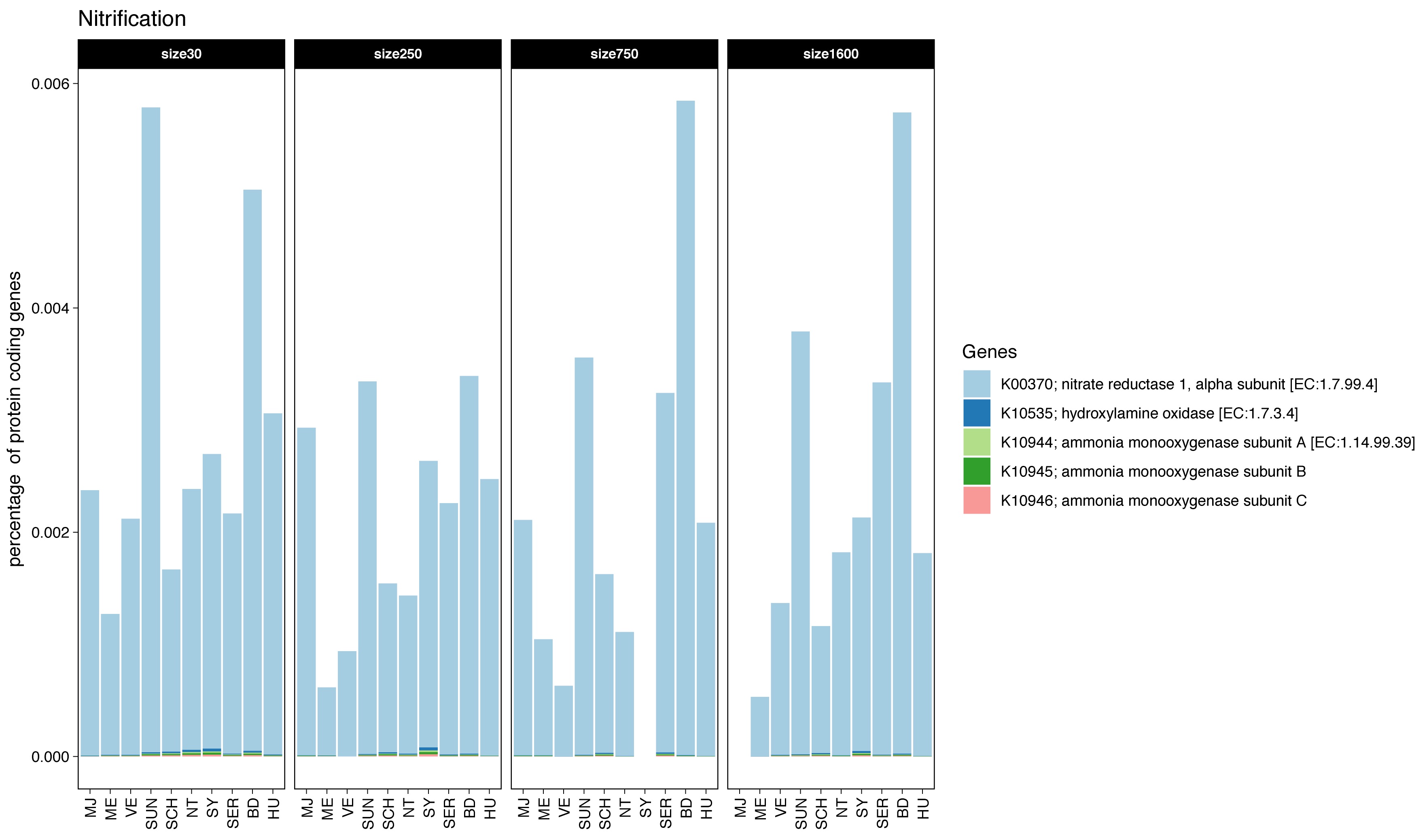

### Supplemental Figure 5

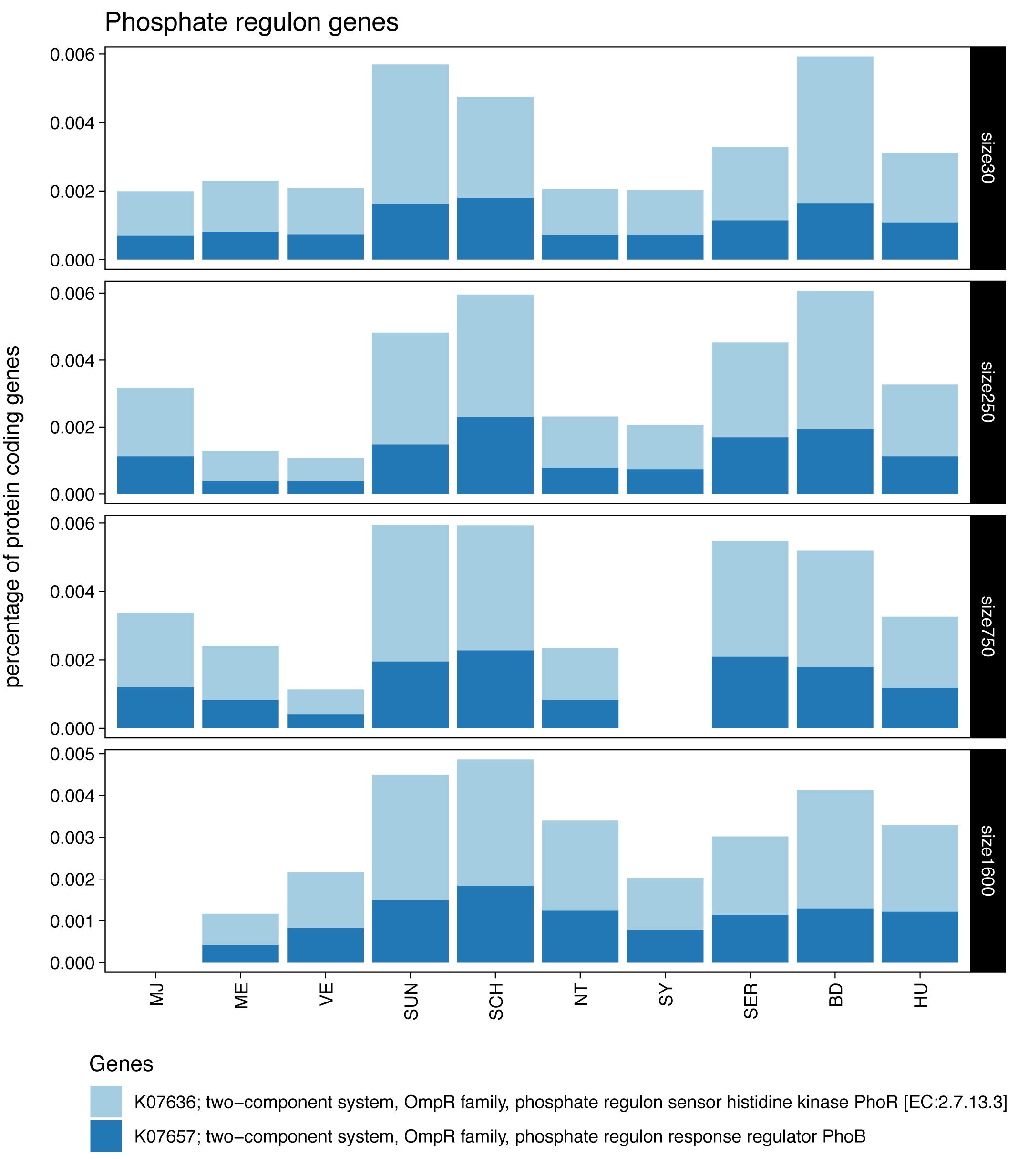
