## Supplemental Figure caption for "Bacterial community changes with cryoconite granule size and their susceptibility to exogenous nutrients on 10 glaciers in northwestern Greenland"

Supplemental information

SI figure 1: Location of sampling sites on 10 glaciers in Qaanaaq region, Northwestern Greenland. Glacier name with abbreviation and coordinate are listed in SI Table 1.

SI figure 2: Relative abundance of ASVs in major genera of each cryoconite granule size category (Size30, Size250, Size750, Size1600) in 10 glaciers around Qaanaaq. Limited number of ASVs were shown for genus: *Phormidesmis* (relative abundance more than 0.0005), *Acidiphilium* (>0.005), *Granulicella* (>0.003), *Deinococcus* (>0.001) and *Leptolyngbya* (>0.003).

SI figure 3: Principal coordinates plots of all samples from 10 glaciers based on Bray-Curtis distances. Plots are separated by size category of cryoconite granule.

SI figure 4: Prediction of the abundance of nitrification related genes (nitrate reductase, Hao, amoA, amoB, amoC) based on community structure. Abundance of functional genes was predicted based on the bacterial community structure using Tax4Fun.

SI figure 5: Prediction of the abundance of phosphate regulon related genes (PhoR, PhoB) based on community structure. Abundance of functional genes was predicted based on the bacterial community structure using Tax4Fun.

SI Table 1: Information about sampling sites on 10 glaciers.

SI Table 2: List of genera with higher similarity percentage (SIMPER > 1.0) in three categories (1: All size is for genera have higher similarity throughout all sizes, 2: Small size is for genera have higher similarity only in smaller granules, and 3: others is for genera was excluded with above 2 categories)

SI Table 3: List of genera, which are estimated as indicator genera for each granule size using indicator species analysis (indval function of “labdsv” packages of R).

SI Table 4: Results of PERMANOVA tests on the effects of nutrients (ammonium, nitrate, phosphate, silicate) on the Bray Curtis and weighted UniFrac distances.

changes between samples

by analysis of similarities (ANOSIM) from two beta diversity methods Bray Curtis, Weighted UniFrac) changes by changes by granule size changes

R
